# CAF contractility controls collagen stiffness through history-dependent competition between mechanical and proteolytic remodeling

**DOI:** 10.64898/2026.09.19.752881

**Authors:** Bashar Emon, Ahmadreza Kashefi, M. Taher A. Saif

## Abstract

Mechanical remodeling by cancer-associated fibroblasts (CAFs) stiffens tumors and can restrict molecular transport, yet the time-resolved coupling between cellular force and matrix mechanical state is poorly defined. Here we use a microfabricated force sensor integrated with 3D CAF-collagen tissue to measure CAF-generated force and collagen stiffness continuously during pharmacological perturbation. ROCK or non-muscle myosin II inhibition reduced force and remarkably drove collagen stiffness below its initial value rather than merely arresting stiffening. This softening was prevented by broad-spectrum MMP inhibition, but only modestly altered by lysyl oxidase inhibition, supporting a competition between force-driven collagen compaction and proteolytic remodeling. Second-harmonic generation imaging showed shorter, thinner and less aligned fibers after ROCK inhibition, while fluorescence recovery after photobleaching indicated faster dextran transport. Moreover, CAF contractile adaptation depended strongly on the duration of inhibition, with reversible recovery after brief treatment, a hyper-contractile rebound following prolonged treatment and washout, and partial force recovery during continuous exposure. These findings identify CAF-matrix mechanics as a dynamic, treatment-history-dependent balance and provide a framework for designing stromal interventions that alter stiffness and transport.

## Introduction

The mechanical state of a solid tumor is an active regulator of disease progression rather than a passive consequence of growth [1–4]. Increased stromal stiffness, now recognized as a physical hallmark of many solid tumors [5], promotes invasion [6,7], activates mechanosensitive pathways including YAP/TAZ [8–10], and can impede molecular transport and therapeutic delivery [4,11]. Cancer-associated fibroblasts (CAFs) are major drivers of this mechanical remodeling [12]. Through RhoA-ROCK-dependent actomyosin contractility [10,13], CAFs compact, align and tension collagen networks [6,14], promote LOX-dependent matrix crosslinking [15,16], and progressively stiffen the tumor microenvironment over disease-relevant timescales (Fig. 1A) [2,17,18].

**Figure 1.**
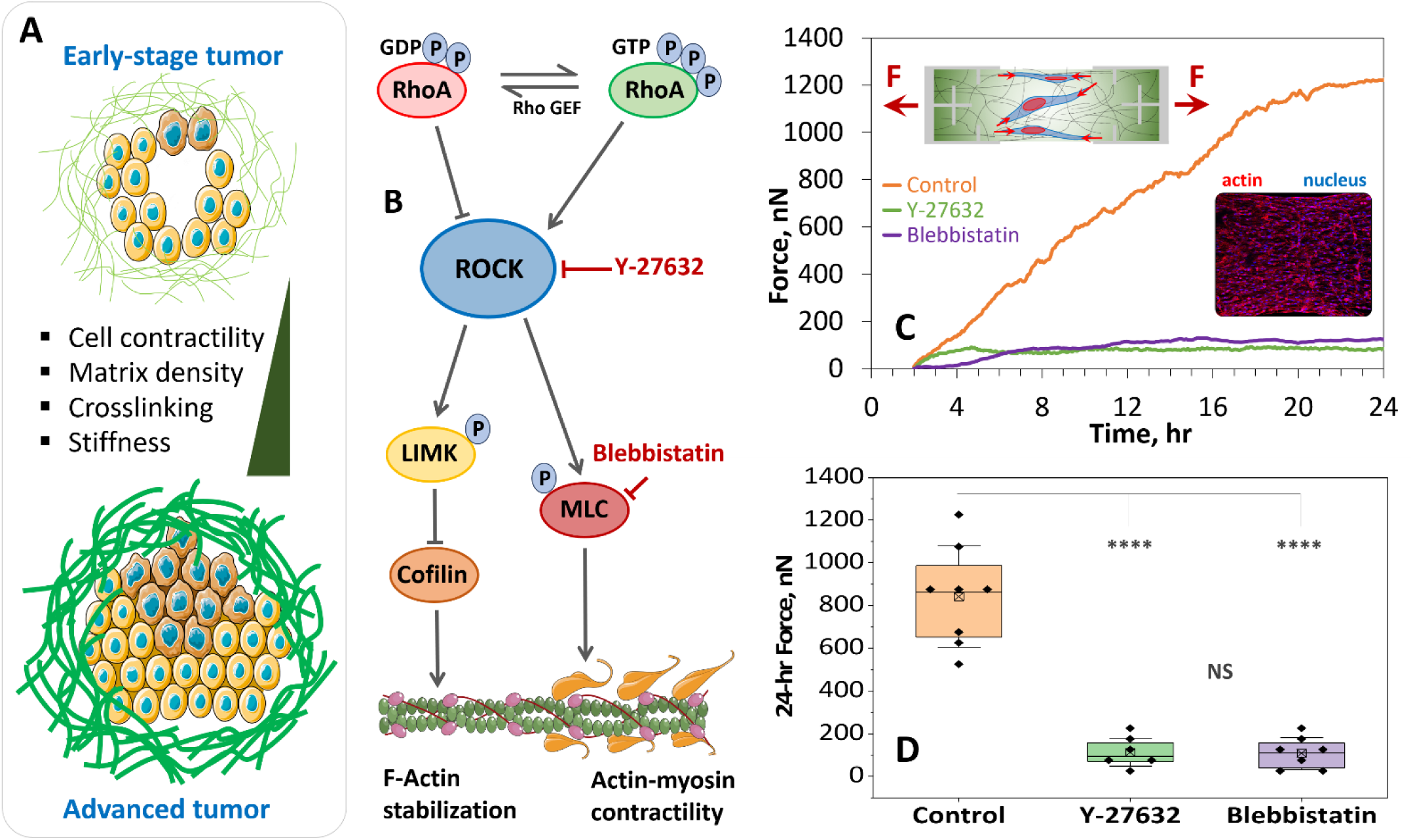
RhoA–ROCK–myosin signaling regulates force generation capacity in CAFs. (A) Schematic illustrating extracellular matrix (ECM) remodeling during tumor progression. Early-stage tumors are characterized by lower cell contractility and reduced matrix density, crosslinking, and stiffness, whereas advanced tumors display increased cellular contractility and a denser, stiffer ECM. (B) Diagram of the RhoA-ROCK signaling pathway regulating actin cytoskeletal dynamics and actomyosin contractility. Activation of RhoA stimulates ROCK, which phosphorylates LIMK and myosin light chain (MLC), promoting F-actin stabilization and actin-myosin contractility. Pharmacological inhibition of ROCK (Y-27632) or myosin II (Blebbistatin) disrupts force generation [37,38]. (C) Representative time courses of tissue force measured over 24 hours for control cells and cells treated with Y-27632 or Blebbistatin, showing reduced force development upon pathway inhibition. Insets: schematic of the force measurement setup illustrating cell-generated forces acting within 3D collagen ECM. This force is transmitted to the sensor and recorded. Representative fluorescence image of the CAF-collagen tissue stained for F-actin (red) and nuclei (blue), demonstrating cellular organization within the CAF-collagen tissue. (D) Quantification of total contractile force at 24 hours. Both Y-27632 and Blebbistatin significantly reduce force generation compared with control, with no significant difference between inhibitor treatments. Box plot – 25th, median, 75th percentile; x-mean; whiskers - SD. Statistical significance was determined by one-way ANOVA with Tukey’s post hoc testing, **** p<0.0001. Illustration adapted from SMART [39]

These observations have motivated strategies intended to normalize the tumor stroma by altering CAF activation or extracellular-matrix composition [12,19]. Approaches targeting TGF-beta signaling [20–22], LOX-dependent collagen crosslinking [23–25] and hyaluronan [26,27] have shown preclinical promise, but clinical translation has been inconsistent. A central unresolved issue is therefore not simply whether stromal mechanics can be perturbed, but how CAF-generated force, matrix structure and enzymatic turnover evolve together during and after treatment.

Against this backdrop, ROCK inhibition can attenuate collagen deposition and improve enhance drug uptake in some cases [28–30]. However, such endpoint results lack critical insights on whether reduced contractility merely removes an active tensile contribution to tissue stiffness, arrests further remodeling or shifts the matrix toward an alternative remodeling trajectory. Nor is it known whether CAF force remains suppressed after drug withdrawal or adapts during sustained inhibition. Addressing these questions requires simultaneous, longitudinal measurements of cell-generated force and matrix mechanical state in a three-dimensional microenvironment.

Here we use a microfabricated sensor platform [31,32]that measures tissue force and stiffness in the same CAF-collagen tissue over time. We combine contractility perturbations with MMP and LOX inhibition, second-harmonic generation (SHG) imaging and fluorescence recovery after photobleaching (FRAP). The data supports a model in which collagen mechanics emerge from competition between force-driven compaction and proteolytic remodeling and show that recovery from ROCK inhibition depends strongly on treatment history. These results define a dynamic force-stiffness relationship relevant to the timing and durability of stromal interventions.

## Results

### Microfabricated sensor enables longitudinal force and stiffness measurements in 3D CAF-collagen tissues

To determine how CAF-generated force governs the evolving mechanical state of the surrounding matrix, we first established a 3D tissue construct that enables longitudinal measurement of both cellular force and tissue stiffness. CAF-populated collagen I tissue specimens were formed directly between two grips of a microfabricated PDMS sensor (Fig. S1A). One grip was coupled to a compliant sensing spring, allowing cell-generated tissue tension to be quantified from the calibrated spring constant and optically measured displacement [31,32]. Brightfield and confocal imaging confirmed formation of a continuous tissue spanning the grips, with cells distributed throughout the tissue thickness (Fig. S1B-E).

The same device was also used for longitudinal mechanical testing by applying controlled axial deformation and measuring the corresponding force response. Tissue stiffness was determined from the initial linear slope of the force-displacement curve. This configuration therefore enabled repeated measurement of both spontaneous CAF-generated force and tissue stiffness in the same tissue over time, allowing changes in active contractility to be distinguished from persistent remodeling of matrix mechanical properties.

### ROCK and myosin II inhibition suppress CAF force in 3D collagen

Non-muscle myosin II (NM-II) is the principal actomyosin motor that generates fibroblast contractile force [33–36]. RhoA activates ROCK, which promotes myosin light-chain phosphorylation and stabilizes F-actin through LIM kinase and cofilin signaling (Fig. 1B).

We therefore perturbed this pathway at two levels: Y-27632 inhibited ROCK upstream of NM-II, whereas Blebbistatin directly inhibited NM-II ATPase activity.

Based on this framework, we hypothesized that CAF-generated contractility drives collagen tissue stiffening and that suppressing force generation would reduce this mechanical remodeling. Using inhibitors at two distinct points in the contractile pathway allowed us to test whether this effect was specific to ROCK signaling or reflected a more general dependence on actomyosin-generated force. CAFs were embedded in 2 mg/mL collagen between the sensor grips, and force was monitored continuously for 24 h during treatment with 10 μM Y-27632 or 20 μM Blebbistatin.

Untreated tissues developed force progressively, reaching approximately 700-1000 nN by 24 h (Fig. 1C). In contrast, tissues treated with either Y-27632 or Blebbistatin shortly after tissue formation (t ∼ 4-5 h) plateaued near 100 nN force, indicating sustained suppression of force development over the subsequent hours. At 24 h, both inhibitors significantly reduced tissue force relative to untreated controls, with no detectable difference between Y-27632 and Blebbistatin (Fig. 1D). The comparable effects of upstream ROCK inhibition and direct NM-II inhibition indicate thattissue force is predominantly actomyosin-dependent.

### ROCK inhibition produces history-dependent recovery

CAF-generated mechanical force contributes to matrix remodeling, tissue stiffening, tumor invasion and therapeutic resistance [1,2,5,10,22,24,40–42]. Because ROCK inhibitors are being considered as anti-fibrotic and anti-stromal strategies [43–47], we next asked whether suppression of CAF force was reversible and whether recovery depended on treatment duration. Three Y-27632 regimens were compared (Fig. 2A-C): a brief approximately 2 h exposure followed by washout (control regimen, Fig. 2A); continuous exposure during the first 22 h followed by a second approximately 2 h exposure (∼24 h total) and washout (D1_Y27, Fig. 2B); and drug addition after 24 h followed by continuous exposure without washout (D2_Y27, Fig. 2C). These regimens separated acute reversibility, recovery after prolonged pretreatment and adaptation during sustained inhibition.

**Figure 2:**
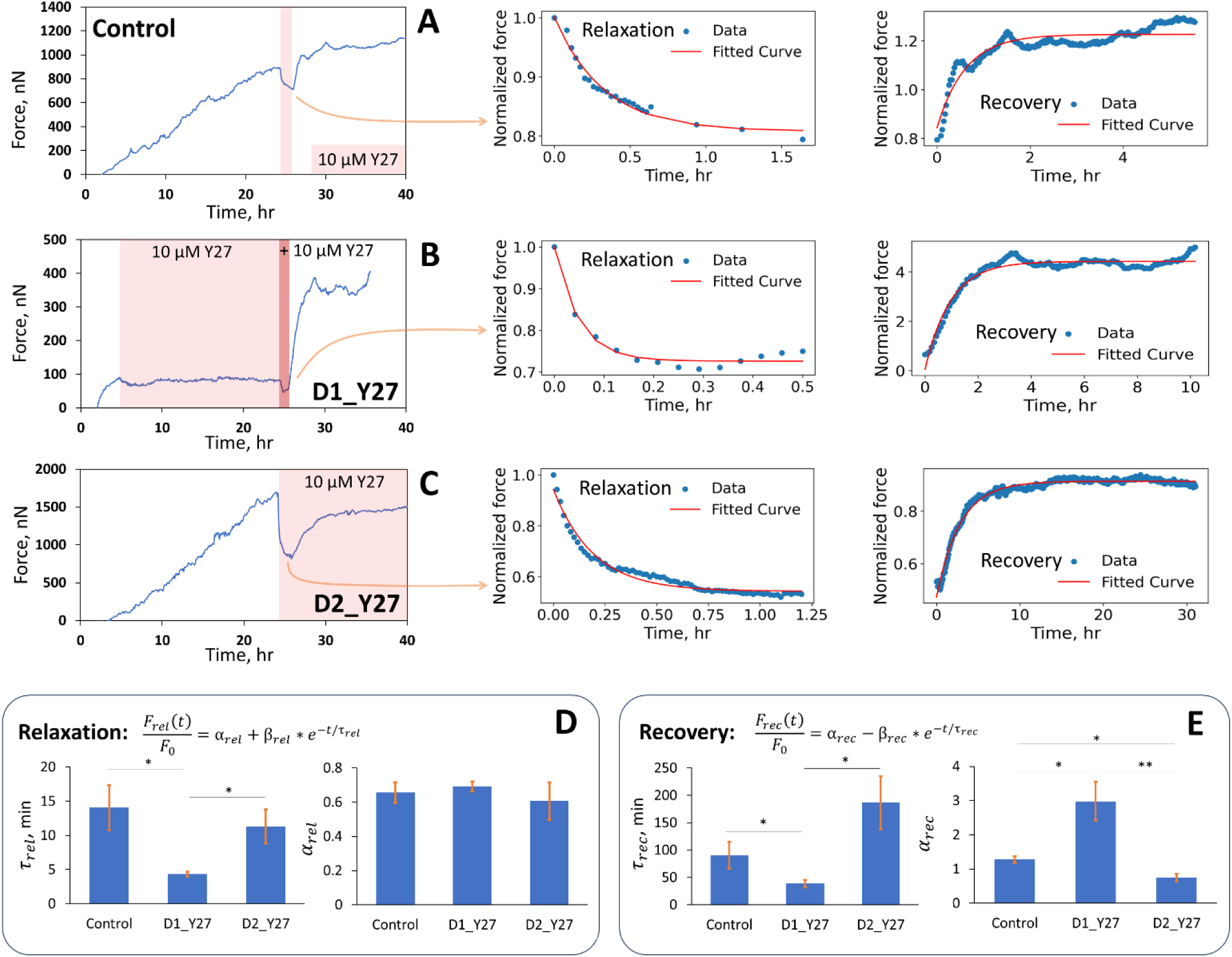
Quantification of force relaxation and recovery dynamics following inhibition of Rho-kinase (ROCK). (A-C) Representative time-lapse force traces showing CAF contractility before, during, and after treatment with the ROCK inhibitor Y-27632 (Y27) that reduces non-muscle myosin II (NM-II) activity. (A) Control tissues were monitored continuously for 24 hours without perturbation, followed by 2 hours of Y-27632 treatment, drug washout, and an additional 24-hour observation period. **(B)** D1_Y27 condition showing force evolution during continuous Y-27632 exposure during the first 24 hours after tissue formation, followed by an additional drug application for ∼2 hours. Force stayed suppressed with continued drug exposure for the first day. Additional dose further reduced force. After washout, the tissue went into a hypercontractile state with force reaching nearly three times the pre-treatment levels. **(C)** D2_Y27 condition showing long-term force dynamics in tissues treated with Y-27632 after 24 hours of culture and subsequently maintained in drug-containing medium for an additional 24 hours. For this condition, we observed limited recovery under sustained drug exposure, suggesting that CAFs adapt to prolonged ROCK inhibition by partially restoring contractile capacity. For relaxation and recovery analyses, raw force traces (left panels) were normalized to the pre-treatment force level, and the relaxation and recovery phases were fit using exponential models (middle and right panels; blue dots represent experimental data, and red lines indicate fitted curves). Relaxation curves quantify the decay in normalized force immediately following Y-27632 administration, while recovery curves capture force redevelopment after drug removal or force evolution during continued drug exposure. **(D-E)** Bar graphs summarize extracted fitting parameters, including relaxation time constants (*τ_rel_*), recovery time constants (*τ_rec_*), minimum normalized force (*α_rel_*), and recovered force amplitude (*α_rec_*) across conditions. Data are presented as mean ± S.E.M. Statistical significance was determined by one-way ANOVA with post hoc testing (*p < 0.05, **p < 0.01).

Force traces were normalized to the value immediately before the analyzed perturbation. Relaxation and recovery phases were fitted with single-exponential functions because actomyosin turnover, focal-adhesion remodeling and cytoskeletal reorganization often produce first-order force responses [48–53]; related exponential descriptions have been used for fibroblast force relaxation and recovery [49,53,54]. The fits provide phenomenological measures of how rapidly force dissipated and redeveloped, without identifying the molecular steps governing either process.

For the relaxation phase (Fig. 2A-C), *τ_rel_* represents the characteristic time required for force dissipation after drug administration, with smaller values indicating faster relaxation (Fig. 2D-E). The parameter *α_rel_* represents the minimum normalized force reached during inhibition and therefore reflects the extent to which contractility can be suppressed (Fig. 2D-E). For the recovery phase (Fig. 2A-C), *τ_rec_* represents the characteristic time required for force redevelopment after washout or during continued drug exposure, with smaller values indicating faster recovery (Fig. 2D-E). Finally, *α_rec_* represents the recovered force amplitude relative to the pre-treatment force level and reflects the degree to which CAFs can restore their contractile capacity (Fig. 2D-E). Together, these parameters provide complementary information on both the kinetics and magnitude of contractile adaptation following chemical intervention (Table 1).

**Table 1:** Parameters from single-exponential fits of force relaxation and recovery after Y-27632 treatment. Values are mean +/- s.d. ***τ_rel_*** and ***τ_rec_*** are the characteristic relaxation and recovery times, respectively; ***α_rel_*** is the minimum normalized force during inhibition; and ***α_rec_*** is the recovered force relative to the pre-treatment value.

| Condition | Treatment regimen | $\tau_{rel}$,<br>min | $\alpha_{rel}$ | $\tau_{rec}$,<br>min | $\alpha_{rec}$ | Interpretation |
| --- | --- | --- | --- | --- | --- | --- |
| **Control** | ~2 h Y-27632 followed by washout | 14.0 ± 7.4 | 0.66 ± 0.13 | 90.5 ± 41.8 | 1.28 ± 0.19 | Reversible force suppression with recovery slightly above baseline |
| **D1_Y27** | 24 h pre-exposure + ~2 h Y-27632 followed by washout | 4.3 ± 0.8 | 0.69 ± 0.06 | 38.6 ± 15.0 | 2.98 ± 1.26 | Faster relaxation and strong hypercontractile rebound after washout |
| **D2_Y27** | Y-27632 added on Day 2 and maintained continuously | 11.3 ± 5.6 | 0.61 ± 0.22 | 186.6 ± 96.4 | 0.75 ± 0.26 | Slow partial force recovery despite continued ROCK inhibition |

In the control tissues (Fig. 2A), brief ROCK inhibition with 10 μM Y-27632 induced a rapid decay (*τ_rel_*∼15 *min*) (Fig. 2D) in force, followed by recovery to pre-treatment levels (*α_rec_*∼1.3) (Fig. 2E) within 2 hours (*τ_rec_*∼90 *min*) (Fig. 2E) after washout, indicating that acute ROCK inhibition was therefore largely reversible over the observation period.

In the samples that were pre-exposed to a day-long treatment of 10 μM Y-27632 (Fig. 2B), an additional dose of 10 μM Y-27632 triggered a faster decline (*τ_rel_*∼4 *min*) (Fig. 2D) in the already suppressed force. Remarkably, complete removal of the drug prompted an extreme surge in contractility, with force jumping to nearly three times the pre-treatment level (*α_rec_*∼3, *τ_rec_*∼40 *min*) (Fig. 2E), indicating a *hyper-contractile* response following prolonged ROCK inhibition. Notably, drug-containing medium was added about ∼4-5 h after the cell-collagen constructs had polymerized; thus, Y-27632 was applied early during cell-mediated tissue compaction, but not during initial collagen gelation.

When 10 μM Y-27632 was added after 24 h and maintained continuously (Fig. 2C), a rapid decline in force with an intermediate relaxation time constant (*τ_rel_*∼11 *min*) was observed (Fig. 2D). Surprisingly, despite the continued presence of the inhibitor, tissues gradually regained force over time, reaching lower than pre-treatment level (*α_rec_*∼.75) (Fig. 2E). This slow (*τ_rec_*∼180 *min*) recovery (Fig. 2E) is consistent with cellular adaptation, incomplete pathway blockade or compensatory force generation, although the responsible mechanism was not resolved. Across regimens, the initial force reduction was similar (*α_rel_*∼.6 -. 7, approximately 60-70%), whereas recovery was strongly treatment-history dependent.

### CAF contractility controls matrix stiffness

CAF-generated forces compact and align collagen, increasing tissue stiffness and altering tumor-cell signaling, invasion and transport [10,24,41,55–60]. We used the sensor to determine whether contractility inhibition simply prevented further stiffening or actively changed the mechanical state of the matrix.

Acellular collagen constructs generated negligible force and showed no measurable change in stiffness after 24 hours polymerization (Fig. 3A). The matrix was therefore mechanically stable over the experimental interval in the absence of cells.

**Figure 3:**
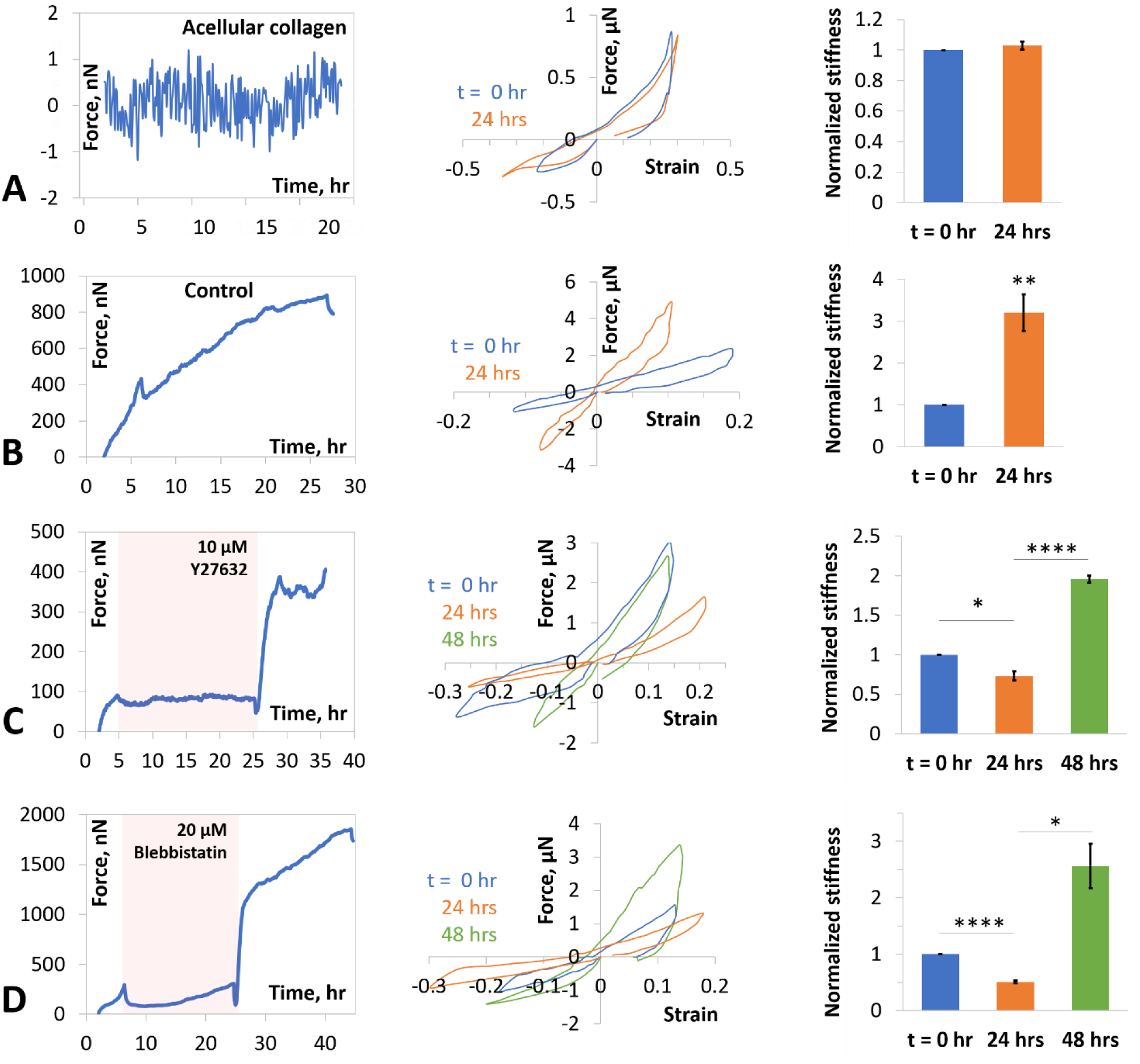
Collagen remodeling by CAFs is force dependent. Force history, force-strain curves from stiffness test and normalized stiffness values for samples **(A)** without cells as ‘acellular control’ **(B)** with CAFs and without drugs as ‘control’, **(C)** with CAFs and Y-27632, and **(D)** with CAFs and Blebbistatin. Stiffness was measured from the slope of the force-displacement plots and subsequently normalized with respect to the initial stiffness of the same sample. **(A)** Acellular control samples without cells show that collagen does not produce tension and the stiffness of the matrix remains unchanged after polymerization. **(B)** CAFs, without exposure to any drug, remodel collagen within 24 hours and increase stiffness by ∼3 folds. **(C-D)** Force inhibition by Y-27632 and Blebbistatin results in a reduction of stiffness of the matrix in 24 hours. After removal of the drugs, CAFs demonstrate hypercontractile recovery and stiffen the matrix in the following 24 hours. Data are presented as mean ± S.E.M. Statistical significance was determined with T-test (*p < 0.05, **p < 0.01, ***p < 0.001, ****p < 0.0001).

Untreated CAF-containing tissues developed progressively increasing force and became more than threefold stiffer over 24 h (Fig. 3B). This contrasted with the acellular controls and demonstrated that CAF activity, rather than passive collagen aging, drove the measured reinforcement.

To determine whether controlling cell contractility can attenuate matrix stiffening, CAFs were treated with either Y-27632 or Blebbistatin during the 24-hour culture period (Fig. 3C-D). We initially expected that suppressing CAF force generation would largely prevent active matrix remodeling, resulting in minimal change collagen stiffness. Surprisingly, however, inhibition of contractility not only prevented matrix stiffening but reduced collagen stiffness below the initial baseline level, which ranged from 100 to 200 kPa. This softening response was observed with both Y-27632 and Blebbistatin treatment, indicating that suppression of the ROCK/NM-II pathway shifts CAF-mediated remodeling from matrix stiffening toward matrix softening.

Time-resolved stiffness measurements after Y-27632 addition separated the immediate mechanical effect of force loss from subsequent matrix remodeling (Fig. S2). Initially, progressively increasing forces stiffened the matrix over the first 24 hours. However, shortly after Y-27632 addition, tissue force rapidly decreased and was accompanied by a marked reduction in stiffness. This indicates the direct contribution of active contractility to mechanical stiffness of the tissue. Prolonged treatment further reduced matrix stiffness, indicating that the softening response emerges dynamically following suppression of contractility rather than simply reflecting a drop in force (Fig. S2). The delayed component therefore cannot be explained solely by instantaneous unloading and suggests continued structural or enzymatic remodeling of collagen.

After inhibitor (e.g. Y-27632 or Blebbistatin) removal, CAF force recovered andthe tissues re-stiffenedduring the following 24 h (Fig. 3C,D). Matrix stiffness tracked the contractile state of the cells and remained reversible over the duration of these experiments.

### Low force shifts remodeling toward proteolysis

We investigated whether enzymatic collagen turnover contributes to softening after force inhibition. We hypothesized that matrix metalloproteinases (MMPs) and lysyl oxidase (LOX) play a key role in this process, because MMPs can degrade collagen and LOX can stabilize collagen through crosslinking. Our selection of MMP and LOX as candidate regulators was motivated by our previous findings that matrix-remodeling genes are highly mechanosensitive in CAFs [61]. In this study, CAFs cultured on increasingly stiff 2D polyacrylamide gels (1, 10, and 40 kPa) exhibited progressively higher forces (Fig. S3) and corresponding shifts in matrix-remodeling gene expression (Table S1). Specifically, increasing substrate stiffness and cell force were associated with reduced expression of MMP-related genes and increased expression of LOX-related genes (also corroborated by Howard et al [62]).

We next measured secreted MMPs and tissue inhibitors of metalloproteinases (TIMPs) in conditioned medium from CAFs on 1 and 40 kPa gels (Fig. S4). CAFs on the softer 1 kPa substrate (low CAF contractility) secreted significantly more MMP-1 than CAFs on the stiffer 40 kPa substrate (high CAF contractility), whereas TIMP-1 secretion was significantly lower on the softer substrate. Although the remaining measured MMPs and TIMPs did not individually reach statistical significance, they showed a consistent force-dependent trend across the panel: CAFs under lower-force conditions generally secreted higher levels of MMPs and lower levels of TIMPs than CAFs under higher-force conditions. This coordinated response is biologically important because reduced TIMP secretion would lessen endogenous inhibition of MMP activity, thereby amplifying the functional effect of increased MMP secretion. Thus, lower CAF force appears to shift the MMP-TIMP balance toward greater matrix proteolysis, whereas higher CAF force favors reduced proteolytic activity. Together with the transcriptomic data, these findings support the hypothesis that force-dependent regulation of ECM degradation and crosslinking pathways contributes to the matrix softening and stiffening observed in 3D collagen, motivating us to directly test whether inhibition of MMP alters the stiffness trajectory during suppression of CAF contractility (Fig. 4).

**Figure 4:**
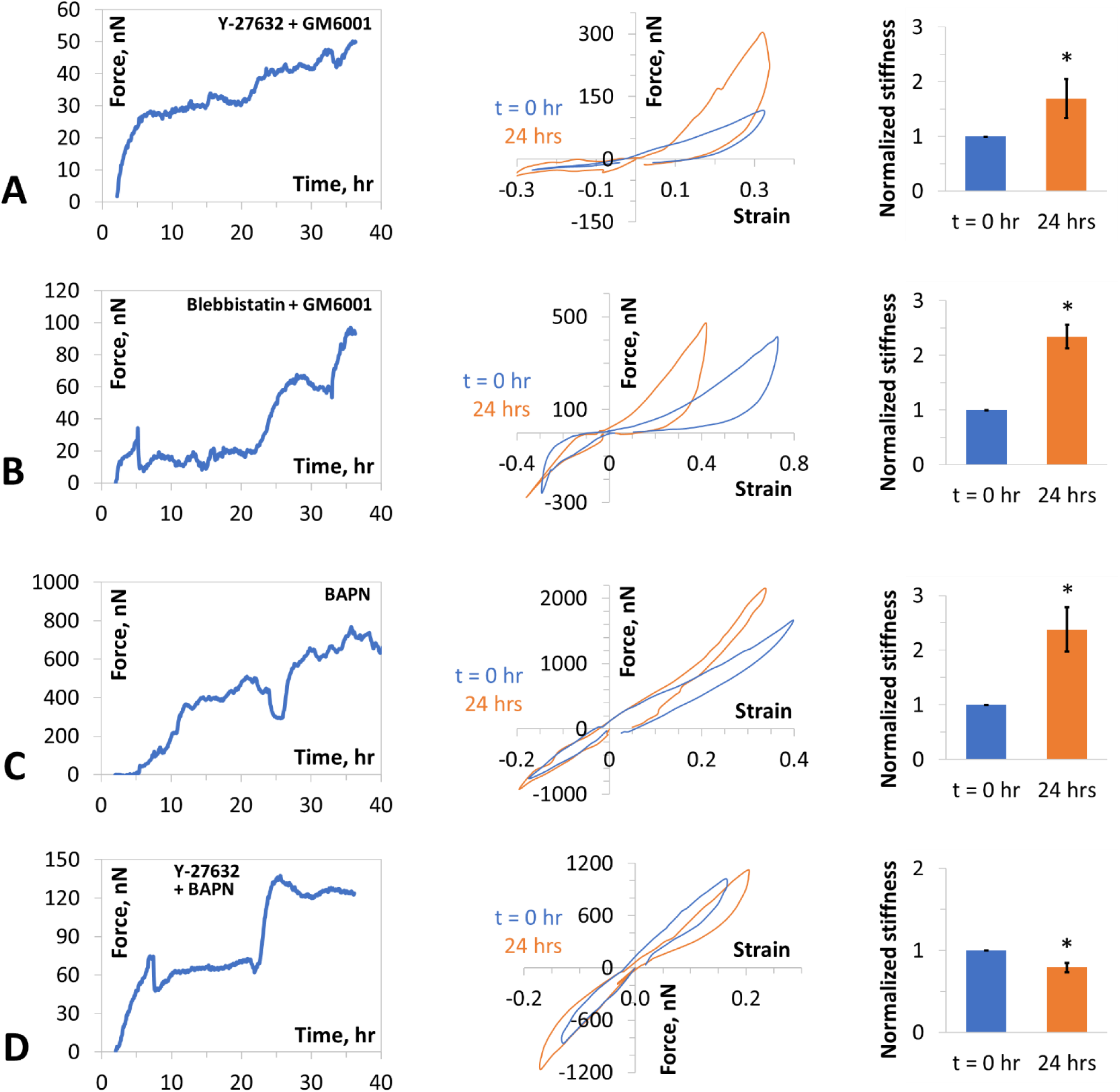
Role of MMP and LOX enzymes in collagen remodeling. Force history, force-strain curves from stiffness test and normalized stiffness for samples with CAFs and **(A)** Y-27632 + GM6001 **(B)** Blebbistatin + GM6001 **(C)** BAPN, and **(D)** Y-27632 + BAPN. Stiffness was measured from the slope of the force-displacement plots and subsequently normalized with respect to the initial stiffness of the same sample. GM6001 is a broad-spectrum MMP-inhibitor, and BAPN is a LOX-inhibitor. MMP inhibition (GM6001, 30 μM) abrogates matrix softening induced by low contractility. LOX inhibition (BAPN, 40 μg/mL) mildly reduces matrix stiffening, indicating cell force induced (plastic) deformation is the primary contributor of collagen stiffening. Column plots: mean ± S.E.M. Statistical significance was determined with T-test (*p < 0.05).

We therefore tested the mechanism directly in the 3D CAF-collagen tissues. Co-treatment with the broad-spectrum MMP inhibitor GM6001 (30 μM) prevented the softening caused by either Y-27632 or blebbistatin (Fig. 4A,B). Stiffness instead increased over 24 h, although less than in untreated controls. This pharmacological rescue indicates that MMP activity is required for net softening when actomyosin force is suppressed.

Inhibition of LOX with beta-aminopropionitrile (BAPN) produced a comparatively modest effect. BAPN alone (40 μg/mL) did not measurably alter tissue force and only partially reduced the increase in stiffness at 24 h (Fig. 4C), suggesting that LOX-dependent crosslinking contributes to matrix stiffening but is not its primary determinant over this experimental period. Combining BAPN with Y-27632 produced force and stiffness responses comparable to Y-27632 alone (Fig. 4D), indicating that LOX inhibition provided no additional mechanical effect once ROCK-dependent contractility was suppressed. Together, these findings support actomyosin contractility, rather than LOX-dependent crosslinking, as the dominant regulator of the observed mechanical response under these conditions.

Together, the perturbations support a competition between mechanical reinforcement and enzymatic degradation. At high force, collagen compaction, alignment and plastic reorganization dominate the measured stiffness. When force is suppressed, MMP-dependent remodeling becomes sufficient to drive the matrix below its initial stiffness. The persistence of substantial stiffening under LOX inhibition suggests that force-driven collagen compaction and plastic deformation, rather than LOX-mediated crosslinking, are the principal contributors to CAF-mediated matrix stiffening. The persistence of these matrix changes after suppression of cellular force is consistent with cell-mediated collagen alignment producing a residual, plastic-like remodeling state rather than a purely elastic deformation. Although these measurements do not establish irreversible plastic strain, they suggest that active cellular remodeling may generate structural changes that are not reproduced by transient mechanical stretching alone.

### ROCK inhibition reorganizes collagen and diffusion

To further elucidate the structural basis of force-dependent matrix remodeling, we performed second harmonic generation (SHG) imaging on collagen matrices remodeled by CAFs with or without Y-27632 treatment (Fig. 5A,C). Quantitative analysis of SHG images with CT-FIRE [63] fiber segmentation revealed that Y-27632-mediated ROCK inhibition substantially reorganized the collagen architecture over 24 hours (Fig. 5B,D). Control CAF-embedded matrices exhibited longer, thicker, and straighter fibers with orientations strongly anisotropic along the axis of boundary constraint (Fig. 5E-H), consistent with active traction-driven fiber alignment, lateral bundling, and tensional straightening mediated by the contractile axis. Matrices with force-inhibited CAFs showed a statistically significant shift toward shorter, thinner, and more curved fibers (Fig. 5E-H), with the straightness index (*d_end-to-end_*/*L_fiber_*) distribution shifting away from unity. This is mechanically consistent with the loss of actomyosin-generated tensile forces, since fibers are no longer held taut by cellular traction, allowing fibers to relax into higher-curvature, lower-tension configurations. The concurrent reduction in fiber diameter and length under ROCK inhibition suggests MMP-mediated matrix degradation. This is consistent with increased accessibility of cleavage sites in tension-relieved, buckled fibers, as mechanical strain is known to suppress collagenolysis by stabilizing the triple-helical structure against local unfolding at MMP recognition sites [64,65].

**Figure 5.**
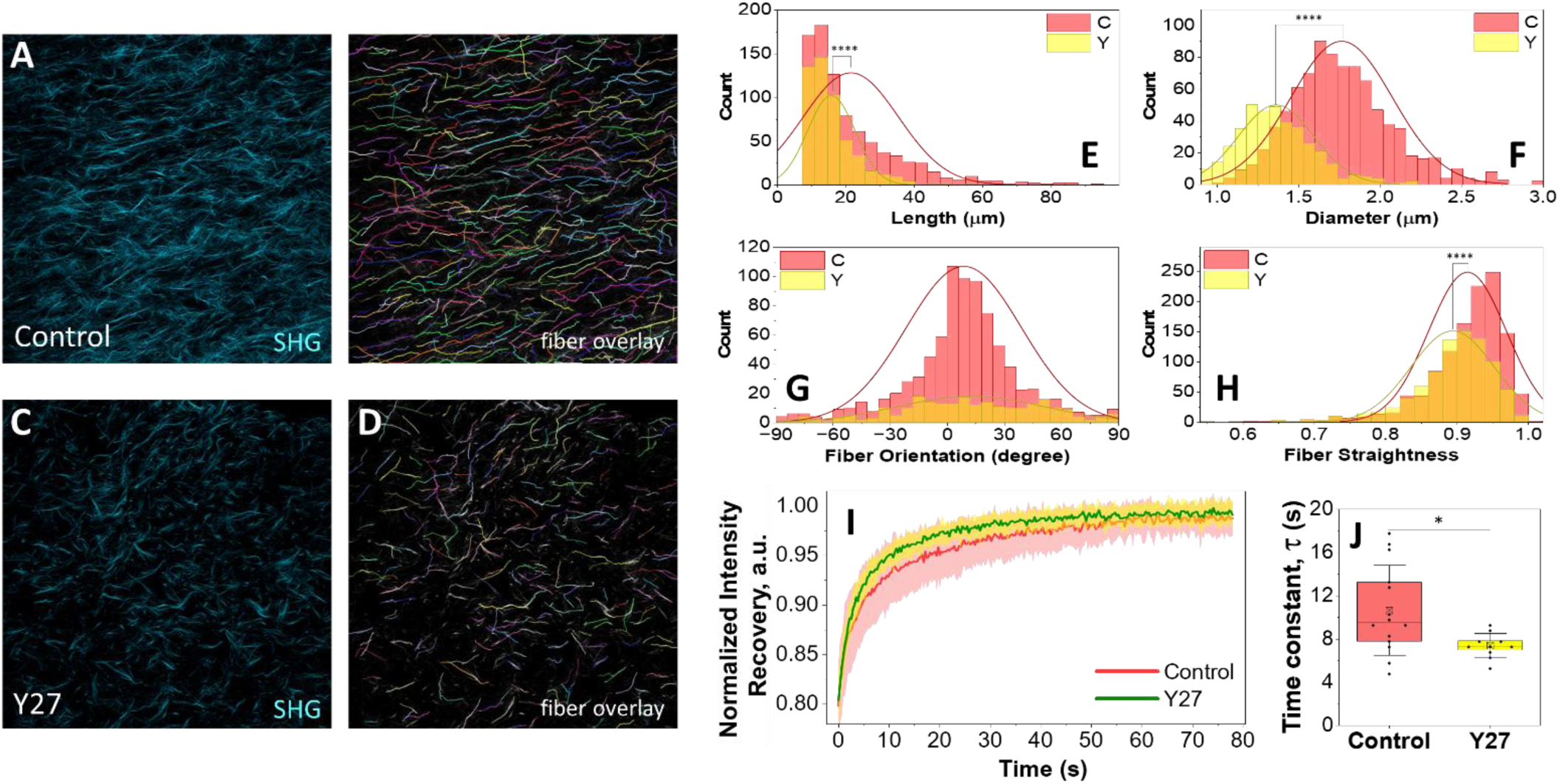
Contractility inhibition alters collagen architecture and transport properties. (A-. **D)** Representative second harmonic generation (SHG) images and corresponding fiber overlays of collagen matrices with **(A-B)** CAFs without drugs (control, C) and **(C-D)** CAFs under 24-hour Y-27632 treatment (D1_Y27, Y). Tissues are oriented left to right along the axis of boundary constraint. Lower SHG intensity per unit area in Y-27632-treated tissues relative to control suggest that ROCK inhibition attenuates contractility-mediated compaction and densification of collagen. Collagen fiber segmentation and reconstruction were performed using CT-FIRE [63]. **(E-H)** Quantitative analysis of collagen fiber morphology and organization. Histograms with overlaid normal distribution fits showing fiber **(E)** length, **(F)** diameter, **(G)** orientation, and **(H)** straightness. Fiber straightness is defined as the ratio of the end-to-end distance to the total fiber length(*straightness* = *d_end-to-end_*/*L_fiber_*), where values approaching 1 indicate straighter fibers. **(I)** Fluorescence recovery after photobleaching (FRAP) analysis using fluorescent dextran. Recovery curves represent normalized intensity over time (line ± shade: mean ± SD). **(J)** Time constant of intensity recovery. Y-27632-treated samples exhibit faster recovery (lower time constant), indicating increased effective diffusivity within the matrix. Box plot – 25^th^, median, 75^th^ percentile; x-mean; whiskers - SD. Statistical significance was determined by T-test, **** p<0.0001, * p<0.05.

To evaluate the functional consequences of these architectural changes on matrix transport properties, we performed fluorescence recovery after photobleaching (FRAP) using fluorescent dextran as a diffusional probe. Y-27632-treated tissues exhibited a reduced recovery time constant relative to control samples, indicating an increased effective diffusivity within the matrix (Fig. 5I). This is coherent with the structural alterations in density, alignment and fiber thickness, reducing steric hindrance to macromolecular transport despite reduced fiber straightness. The coupling between contractility-dependent fiber organization and diffusive transport has direct implications for drug penetration and paracrine signaling within CAF-remodeled tumor stroma.

## Discussion

CAFs are increasingly recognized as regulators of tumor mechanics through their ability to generate force and remodel extracellular matrix [1,66–69], and a stiff microenvironment is a physical hallmark of solid tumors [5]. This study establishes a time-resolved link between CAF-generated force, collagen stiffness, matrix architecture and macromolecular transport. The central finding is that suppressing actomyosin contractility does not simply pause stromal stiffening. Instead, it shifts the net remodeling balance sufficiently far toward degradation that collagen stiffness falls below its initial value. The outcome is also history dependent: brief ROCK inhibition is reversible, prolonged pretreatment primes a strong rebound after washout, and continuous exposure permits partial force recovery.

Using simultaneous force and stiffness measurements in a three-dimensional sensor platform [31,32,70], we distinguish two components of the mechanical response. The rapid decrease in stiffness after Y-27632 addition is consistent with removal of active cell-generated tension. Continued softening over the following hours, together with rescue by GM6001, indicates an additional MMP-dependent change in matrix structure. Thus, the measured stiffness reflects both the instantaneous contractile state and the accumulated history of physical and enzymatic remodeling.

The inhibitor experiments support a working model in which force-driven collagen compaction and alignment compete with proteolytic turnover. Untreated CAFs progressively reinforced the matrix, whereas broad suppression of either ROCK or NM-II exposed a degradative program that drove net softening. MMP inhibition reversed the direction of the stiffness trajectory under low-force conditions, while LOX inhibition produced only a modest reduction in stiffening. These data identify MMP activity as necessary for softening in this model but do not establish which MMPs, substrates or activation mechanisms are responsible.

Previous studies established that ROCK signaling and NM-II activity regulate fibroblast contractility, focal-adhesion maturation and stress-fiber organization [71–78]. Here, treatment history altered contractile recovery independently of the initial magnitude of force suppression. The approximately threefold rebound after prolonged pretreatment suggests that sustained ROCK inhibition changes the state of the cells or their matrix such that force generation is enhanced after washout. Partial recovery during continuous exposure likewise indicates adaptation or pathway bypass. Measurements of myosin light-chain phosphorylation, cytoskeletal organization and focal-adhesion dynamics will be needed to distinguish altered drug sensitivity from compensatory signaling or changes in mechanical loading.

These dynamics are relevant to stromal-normalization strategies targeting ROCK or actomyosin activity [79,80]. A regimen that temporarily improves transport could be followed by rapid re-stiffening if contractility rebounds after drug withdrawal. The current reductionist model does not predict an optimal clinical schedule, but it provides quantitative variables - relaxation, recovery and matrix-stiffness trajectories - that can be used to compare candidate regimens.

Structural and transport measurements provide orthogonal support for the mechanical model. ROCK inhibition reduced fiber length, diameter, straightness and alignment, and accelerated dextran recovery. These observations are consistent with reduced compaction and a more transport-permissive network. Mechanical tension can also protect collagen against enzymatic degradation [64,65], but fiber morphology alone does not prove proteolysis, and FRAP with a single dextran size does not capture convection, binding, vascular delivery or the pharmacokinetics of a therapeutic molecule. The transport conclusion should therefore remain restricted to effective diffusion within the 3D CAF-collagen tissue.

Several limitations define the scope of the conclusions. The study used one commercial colorectal CAF line in a collagen-I-only matrix and did not include cancer, immune or vascular cells; donor-to-donor heterogeneity and reciprocal tumor-stroma signaling were therefore not tested. Y-27632, blebbistatin, GM6001 and BAPN are pharmacological probes with off-target or pathway-wide effects, and genetic perturbations would improve mechanistic specificity. Also, direct collagen-cleavage or MMP-activity measurements were not performed in the CAF-collagen tissues. Finally, the experimentstypically span two days rather than the longer timescales of tumor evolution.

In summary, CAF-generated force sets collagen stiffness through a dynamic competition between physical reinforcement and MMP-dependent remodeling. This coupling is reversible but retains a memory of treatment history, and its structural consequences alter macromolecular diffusion. Accounting for both the immediate loss of cellular tension and the slower remodeling response will be important when designing interventions intended to normalize tumor mechanics or improve therapeutic access.

## Materials and Methods

### Fabrication of biomechanical sensors

Following the process described in our previous work [31,32], the biomechanical sensors were prepared by casting PDMS (Sylgard 184, Dow) from microfabricated silicon molds that were patterned and etched to a nominal depth of 200 μm using photolithography and deep reactive ion etching (DRIE; STS Pegasus ICP). PDMS base and curing agent were mixed at a 10:1 (w/w) ratio, dispensed into the molds, and allowed to fill all features and trenches via capillary micro-molding. The PDMS was cured at 60 °C for 12 h, after which the sensors were carefully demolded.

For assembly, a small rectangular glass (#2 cover glass, Corning) was bonded to the bottom of a glass-bottom Petri dish (60-mm diameter; Corning) using uncured PDMS to serve as an elevated platform. Individual sensors were then affixed to the glass platform with uncured PDMS, with 8 sensors arranged per dish.

### Tissue formation

High-concentration rat-tail collagen I (Corning) was prepared on ice by neutralizing the 9.5 mg/mL stock solution with 1N NaOH, 10X PBS, and deionized water according to the manufacturer’s protocol [81], yielding a final concentration of 4 mg/mL at pH 7.2. The tissue precursor solution was then generated by mixing the collagen solution with a cell suspension (4 × 10^6^ cells/mL in culture media) at a 1:1 ratio, resulting in a final collagen concentration of 2 mg/mL and 2 × 10^6^ cells/mL cell density.

The cell-ECM mixture was then pipetted onto the sensor grips to form a capillary bridge. Shortly, the solution filled the grip channels and was allowed to polymerize at 37 °C for 45 minutes. To prevent dehydration, a small volume of media was added to the dish prior to tissue formation. After polymerization, culture media was gently added to the dish to fully immerse the tissues, and the dish was subsequently transferred to the incubator or microscope for experimentation.

### Measurement of tissue force and stiffness

A detailed step-by-step process for measuring cell-generated force andtissue stiffnessis available in our published protocol[32]. Briefly, time-lapse brightfield images of the tissues on sensors were acquired throughout culture using an inverted microscope (Olympus IX81, 20× objective) equipped with environmental control. A motorized stage (Prior Scientific Inc.) enabled automated, multi-location imaging ofmultiple tissuesat defined time intervals.Brightfield images of the tissues were captured with a Neo sCMOS camera (Andor Technology; 323 nm/pixel). Spring displacements (*d_c_*) were determined by analyzing images for grip movement using the template matching plugin in ImageJ software. Tissue force was determined as *F* = *k_s_* ∗ *d_c_*, where *k_s_* = 21.3 *nN*/*μm* for the sensors used for the current study.

For stiffness measurement, the samples were subjected to both tension and compression tests at different time points. Force-displacement curves were plotted, and tissue stiffness was determined from the initial linear slope of the tensile loading curve, corresponding to the small-strain tangent modulus. To account for variability in initial tissue stiffnessacross tissues, stiffness values were normalized to the measurement at the first time point (t = 0 hours), allowing direct comparison of stiffness evolution over time between samples.

### Cell culture

Human primary colorectal tumor cancer associated fibroblasts, CAF05 (Neuromics, Edina, MN, USA), were maintained in VitroPlus III Low Serum, Complete medium (Neuromics, Edina, MN, USA). Cells were grown at 37°C in a humidified incubator with 5% CO_2_.

### Chemicals and Drugs

Y-27632 (Sigma-Aldrich, cat # SCM075) was reconstituted in sterile water to a stock concentration of 10 mM. Blebbistatin (Sigma-Aldrich, cat # B0560) stock solution of 10 mM was prepared in Dimethyl Sulfoxide (DMSO, ATCC). Both stock was stored at -20 °C until use and was diluted in culture medium to the desired working concentrations (i.e. 10 μM for Y-27632 and 20 μM for Blebbistatin) immediately prior to use. Drugs were added directly to the culture media at prescribed time points, and treatments were maintained for the specified durations. All treatments were performed under standard cell culture conditions at 37°C with 5% CO_2_.

MMP-inhibitor GM6001 (Cayman Chemical, cat # 14533), clinically known as Ilomastat or Galardin, was reconstituted in DMSO and diluted to 30 μM in media for application. LOX-inhibitor BAPN (Sigma-Aldrich, cat # A3134) was reconstituted in sterile water for storage and diluted in media to 40 μg/mL for application.

### Polyacrylamide substrate preparation

Polyacrylamide (PA) hydrogels (elastic modulus ∼ 1 and 40 kPa) embedded with fluorescent microspheres (Thermo Fisher Scientific) were prepared as previously described [82–84]. Briefly, gels were polymerized from acrylamide and bis-acrylamide solutions (Sigma-Aldrich), with polymerization initiated using ammonium persulfate (APS; Bio-Rad) and tetramethylethylenediamine (TEMED; Bio-Rad). Fluorescent beads were incorporated near the gel surface to serve as fiducial markers for traction force microscopy [84]. Following polymerization, substrates were functionalized using sulfosuccinimidyl-6-(4′-azido-2′-nitrophenylamino)-hexanoate (Sulfo-SANPAH; Thermo Fisher Scientific) and coated overnight with rat-tail collagen I (Corning) at 25 μg/mL. Cells were subsequently cultured on the functionalized substrates prior to force measurements and conditioned media collection.

### Protein concentration quantification (ELISA)

To quantify matrix metalloproteinase (MMP) and tissue inhibitor of metalloproteinase (TIMP) concentrations in conditioned media from two-dimensional (2D) cultures, cells were seeded onto functionalized polyacrylamide (PA) gels at a density of 25,000 cells per 18-mm circular glass coverslip. After allowing the cells to adhere for 3 h, each coverslip was submerged in 1 mL of culture medium. Two samples were prepared for each PA gel stiffness condition (1 and 40 kPa).

After 48 h of culture, the conditioned medium was collected and centrifuged at 2,000 rpm for 10 min in a refrigerated centrifuge maintained at 4 °C. The resulting supernatants were collected, frozen, and stored at −80 °C until analysis. Samples were subsequently shipped on dry ice to Eve Technologies (Calgary, AB, Canada), where MMP and TIMP concentrations were measured using the Human MMP and TIMP 12-Plex Discovery Assay® Array for Cell Culture and Non-Blood Samples (HMMP/TIMP-C,O).

### Traction force microscopy (TFM)

To quantify cellular traction forces in response to substrate stiffness, fibroblasts were cultured on polyacrylamide (PA) hydrogels embedded with fluorescent microspheres (100 nm diameter, excitation/emission 580/605 nm; Thermo Fisher Scientific) that served as fiducial markers. Preparation and mechanical characterization of the PA substrates were performed as previously described [84]. Fluorescence images of the embedded beads were acquired before and after cell removal to obtain reference and deformed states, respectively. Bead displacement fields were calculated from image pairs and subsequently converted into traction stress maps using established TFM algorithms [85,86]. The total traction force was obtained by integrating traction stresses over the projected cell area. Image processing and displacement analysis were performed using ImageJ and custom analysis routines.

### Imaging and analysis

SHG images were acquired using an LSM 980 confocal microscope (Zeiss) two-photon excitation microscope and 40x water-immersion objective lens. Identical imaging settings (laser power, detector gain, and pixel dwell time) were maintained across all conditions to enable quantitative comparison. For each condition, multiple fields of view and z-stacks were acquired from samples to account for spatial heterogeneity.

Quantitative analysis of collagen fiber morphology and organization was performed using CT-FIRE (Curvelet Transform-Fiber Extraction) software [63,87], developed by the Eliceiri laboratory [88]. SHG images were first preprocessed to enhance fiber contrast and suppress background noise. CT-FIRE applies a curvelet-based transformation to detect curvilinear structures and extract individual collagen fibers from the image. The algorithm segments fibers based on intensity and continuity, followed by reconstruction of fiber centerlines. From the extracted fibers, the following parameters were quantified:

- Fiber length: computed as the arc length of each extracted fiber segment.
- Fiber diameter (width): estimated from the local intensity profile perpendicular to the fiber axis.
- Fiber straightness: defined as the ratio of the end-to-end distance to the total fiber length (*straightness* = *d_end-to-end_*/*L_fiber_*), with values approaching 1 indicating straighter fibers.
- Fiber orientation: calculated as the angle of each fiber relative to a reference axis (aligned with the tissue grips), and used to assess network anisotropy.

Orientation distributions were compiled across all detected fibers, and anisotropy was assessed based on the degree of alignment along the principal axis (0°). All analyses were performed using identical parameter settings in CT-FIRE to ensure consistency across samples. Quantitative outputs were aggregated across multiple images and reported.

Immunofluorescence staining and imaging were performed directly on CAF-collagen I tissues cultured on the biomechanical sensor platform. The in vitro tissues remained attached between the sensor grips throughout fixation, permeabilization, staining, and confocal imaging, thereby preserving the tissue geometry and boundary conditions used during mechanical measurements.

Fluorescence microscopy of the 3D CAF-collagen tissues was performed using an LSM 980 confocal microscope (Zeiss) equipped with 10x and 40x objective lenses.

### Fluorescent Recovery After Photobleaching (FRAP)

FRAP measurements were performed directly on 3D CAF-collagen I tissues formed and cultured on the same biomechanical sensor platform. The sensor-supported tissues remained attached between the grips throughout dextran incubation, photobleaching, and fluorescence-recovery imaging, enabling transport measurements under the same treatment conditions, tissue geometry, and mechanical boundary constraints used for force and stiffness measurements.

We carried out FRAP experiments with LSM 980 confocal microscope (Zeiss) C-Apochromat 40x/1.2W (Zeiss) objective was used for image acquisition. Before the FRAP experiment, the samples were incubated in 70,000 MW Oregon Green^TM^ 488 dextran (Invitrogen Cat#: D7173) for 15 mins. A 488 nm Argon laser was used at 75% power with ∼1 μs exposure per pixel for 30 repetitions. During recovery of the fluorescent signal, images were taken approx. every second. The intensity was tracked and processed in ZEN software (Zeiss). Normalized recovery curves were fitted to the following exponential model to determine time constants for different regions of interest.

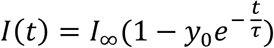

Here, *I*(*t*) is the normalized fluorescence intensity, *I*_∞_ is the normalized plateau intensity, *y*_0_ is the fraction drop after bleach, and *τ* is the time constant of recovery.

### Statistics and reproducibility

For each condition, at least three independent experimental replicates were performed (*n* ≥ 3). All experiments were repeated independently, and consistent results were obtained across replicates. Data is presented as mean ± standard error of the mean (S.E.M.) unless otherwise stated. Statistical analyses were performed using standard parametric tests. For comparisons involving more than two groups, one-way analysis of variance (ANOVA) was used to assess overall differences among group means. When the ANOVA indicated statistical significance, post hoc pairwise comparisons were performed using Tukey’s test. For comparisons between two groups, Student’s *t*-test was applied. A *p*-value of less than 0.05 was considered statistically significant. All statistical analyses and graphical representations were performed using MS Excel and OriginPro 2026.

## Acknowledgement

Research reported in this publication was partially supported by NSF grants ECCS 1934991 and CMMI 2342257, Cancer Center at Illinois (CCIL) seed grant, and the Chan Zuckerberg Biohub Chicago.

## Author contribution

B.E. and M.T.A.S. conceived and designed the study. B.E. and A.K. performed experiments, imaging and analysis. B.E., A.K. and M.T.A.S. prepared the manuscript. All authors have read, edited and approved the final manuscript.

## Data availability

All data needed to evaluate the conclusions in the paper are present in the paper and/or the Supplementary Materials. The authors will make available any additional data/information related to the paper upon request.

## Supplementary Figures

**Figure S1.**
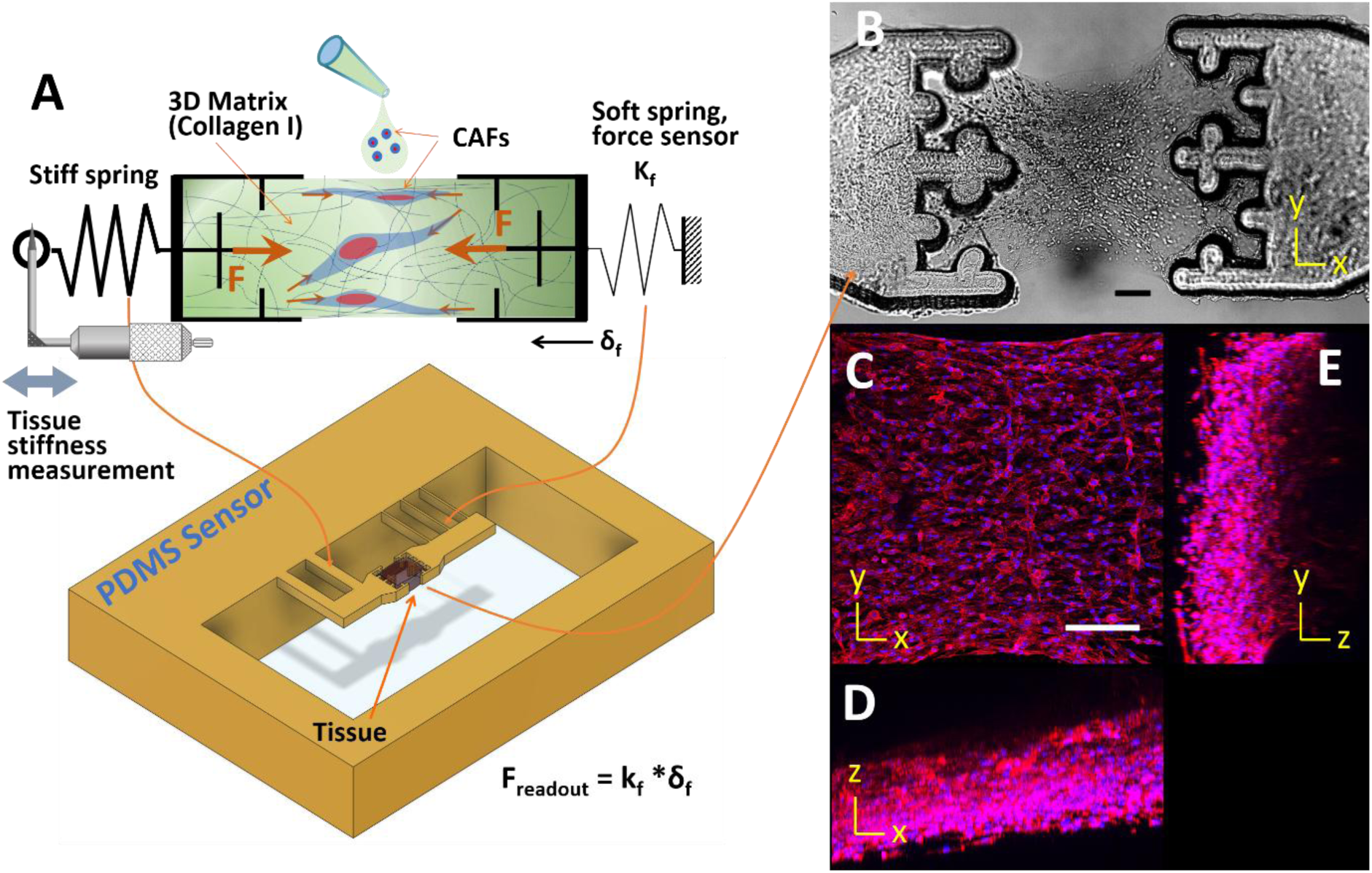
3D CAF tissue integrated with a force-sensing platform. **(A)** Schematic of 3D tissue formation and sensor operation (adapted from [70]). A cell-collagen suspension is introduced between the sensor grips, where it polymerizes and forms a tissue as cells compact and remodel the matrix. Tissue-generated tension is transmitted to the compliant sensing beam, producing a displacement (δ_f_) that is optically tracked and converted to force using its calibrated spring constant (K_f_). Tissue mechanical properties are measured by translating the opposing stiff grip to impose controlled axial deformation while simultaneously recording grip motion and sensing-beam displacement. In the PDMS sensor, narrow beams provide the compliant force-sensing element, whereas wider beams form the stiff loading element. (B) Brightfield image of a 3D CAF-collagen tissue formed between the grips of the force sensor. Cell-generated contractile forces compact the matrix and are transmitted to the sensing elements for continuous force measurement. (C) Confocal fluorescence image of the tissue stained for F-actin (red) and nuclei (blue), revealing extensive cellular spreading and cytoskeletal organization within the collagen matrix (x-y plane). (D,E) Orthogonal x-z and y-z reconstructions of the confocal image stack, respectively, confirming three-dimensional tissue formation and cellular distribution throughout the tissue thickness. Scale bars,200 μm.

**Fig. S2.**
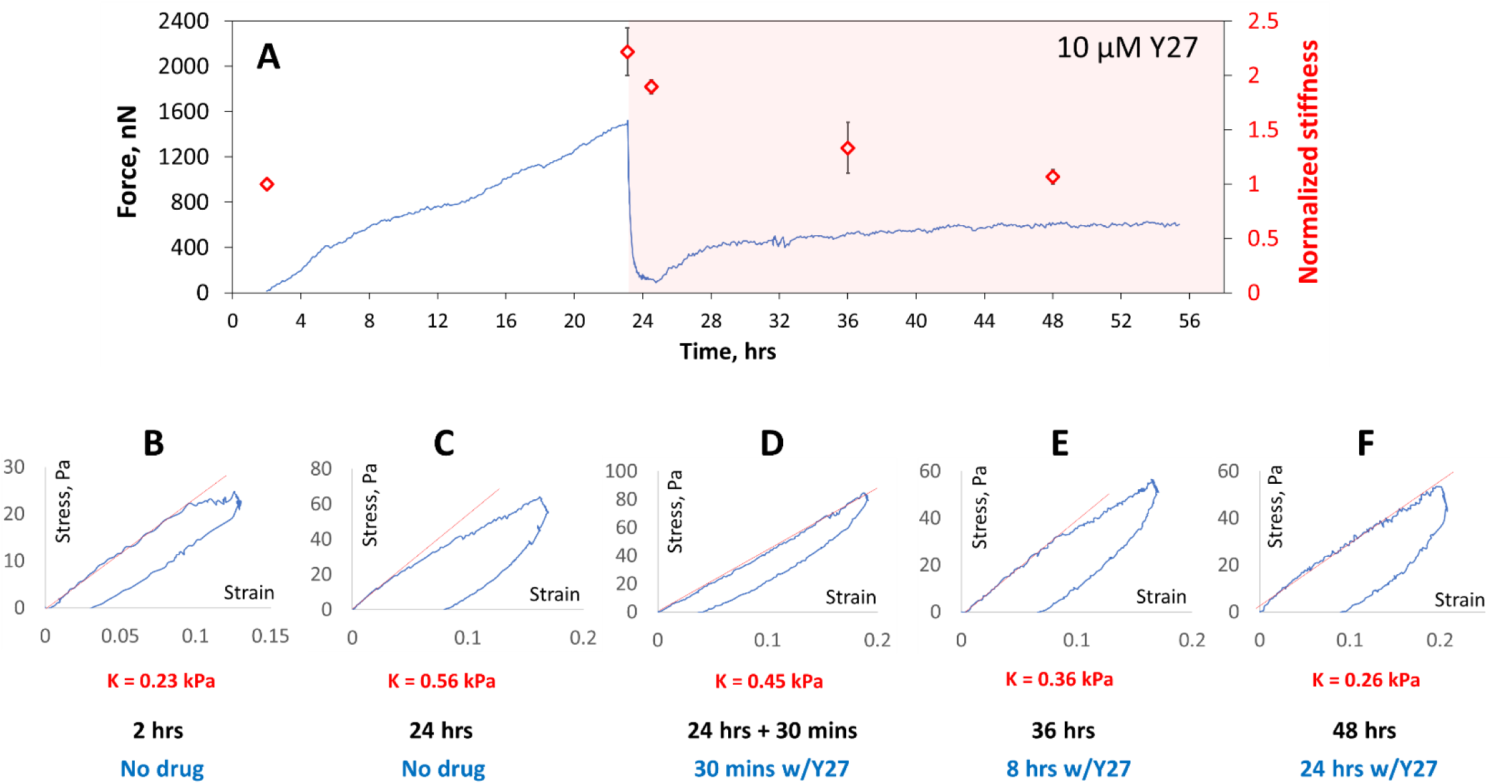
Dynamic changes in CAF force generation and matrix stiffness following inhibition of ROCK-mediated contractility. **(A)** Representative force history showing progressive increase in CAF-generated tension during the first 24 hours of culture, followed by a rapid drop in force immediately after addition of Y-27632. Force partially recovered over time but remained below the pre-treatment maximum. Stiffness history is shown by the data points (red markers) that indicate *mean normalized stiffness* ± *SD*. **(B-F)** Stress-strain curves used to calculate collagen stiffness at different time points. 3D CAF-collagen tissues progressively stiffened the matrix during the first 24 hours in the absence of drug treatment. However, Y-27632 treatment caused an immediate reduction in matrix stiffness within 30 minutes, indicating to the contribution from active contractility. Further softening continued as observed after 12 and 24 hours of treatment. Red lines indicate the initial linear stiffness used for comparison.

**Figure S3.**
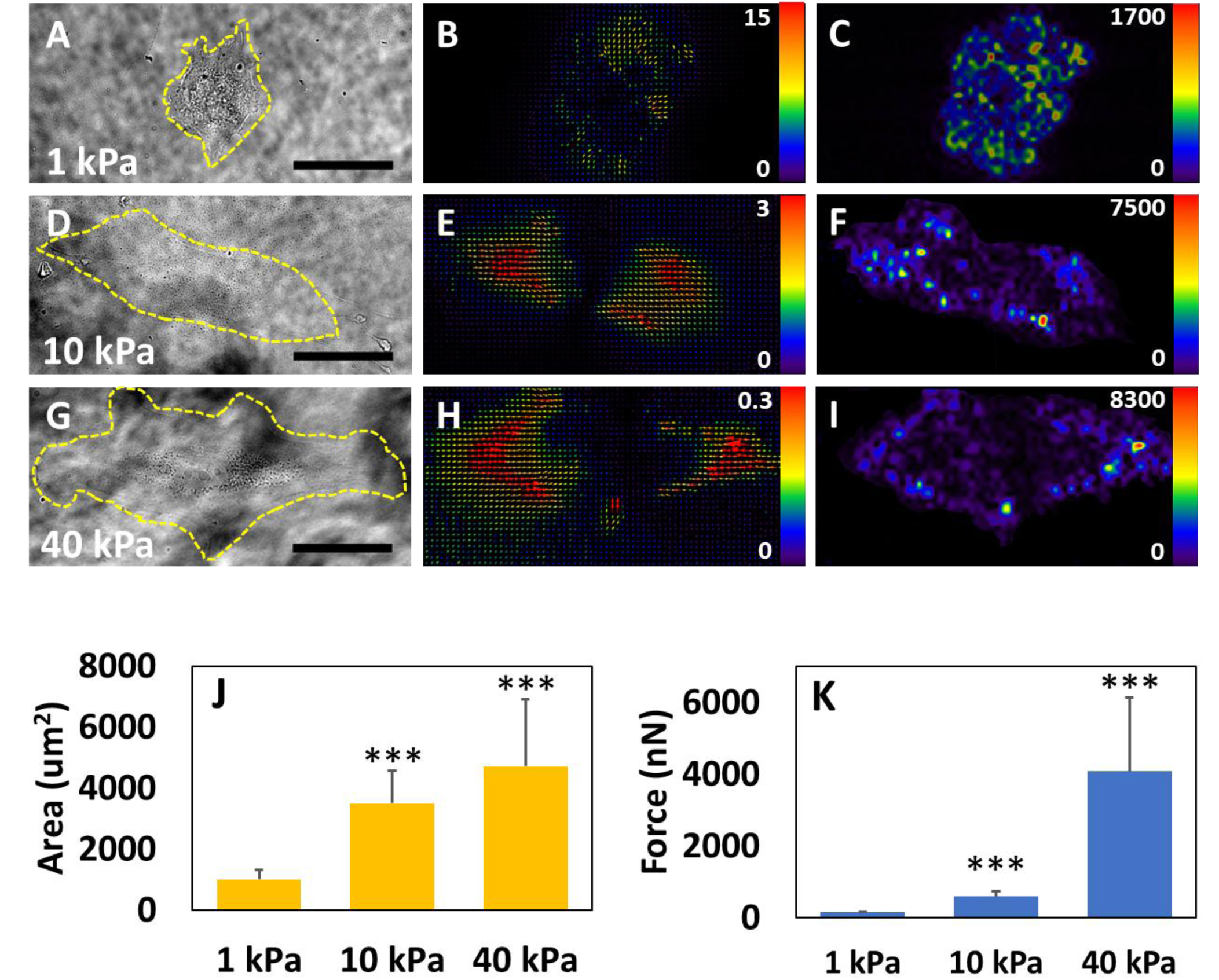
Traction force microscopy results show that CAFs spread more and produce higher force on stiffer substrates. (A-I) Phase contrast, vector displacement field and traction stress heatmap for cells on PA gel substrates with 1 kPa **(A-C)**, 10 kPa **(D-F)** and 40 kPa **(G-I)** stiffness, respectively. **(J-K)** Cell spreading area and total traction force measured from the stress fields (n>10). Error bars: SD. Scale bars: 50 μm. Displacement plot scale unit: μm. Stress map scale unit: Pa. ***p<.0001

**Figure S4.**
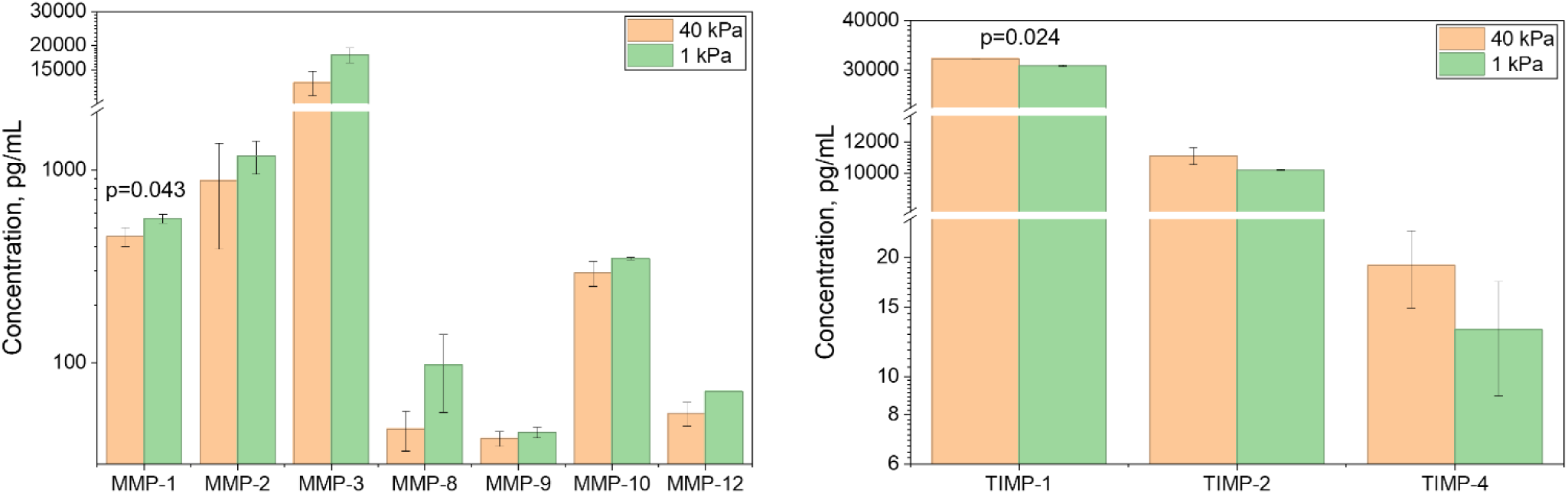
MMP/TIMP secretion profiles of CAFs with different force levels. (cultured on matrices of different stiffness). Concentrations of the measured MMP and TIMP analytes in culture supernatants are shown for cells cultured on 40 kPa substrates (orange) and 1 kPa substrates (green). Bars represent the mean ± SD of two paired replicate measurements (n = 2). K1 and K2 were compared separately for each analyte using a one-sided paired Student’s t-test in the observed direction. MMP-1 and TIMP-1 were significantly different between 40 and 1 kPa (unadjusted p < 0.05); all other analyte pairs were not statistically significant (p ≥ 0.05).

**Table S1.** Statistically significant (p < .05) fold changes (*log_2_ FC,* with respect to 1 kPa) in CAF gene expression relevant to ECM interactions.

|  | <u>1 kPa</u> | <u>10 kPa</u> | <u>40 kPa</u> |
| --- | --- | --- | --- |
| *HAS2* | 0 | 0.6 | 1.6 |
| *SLC8A1* | 0 | 0.8 | 1.6 |
| *PGF* | 0 | -0.1 | -0.9 |
| *ADAMTS7* | 0 | -0.4 | -0.9 |
| *COL3A1* | 0 | -0.5 | -1.0 |
| *COL5A3* | 0 | -0.5 | -1.2 |
| *MMP24* | 0 | -0.1 | -1.2 |
| *LAMA5* | 0 | -0.7 | -1.3 |
| *ITGA10* | 0 | -0.5 | -1.3 |
| *TNXB* | 0 | -0.8 | -1.4 |
| *MMP1* | 0 | 0.1 | -1.4 |
| *BDKRB2* | 0 | -0.6 | -1.4 |
| *VWF* | 0 | -0.3 | -1.4 |
| *ARHGEF6* | 0 | -0.4 | -1.4 |
| *ARHGEF4* | 0 | 0.2 | -1.7 |
| *LAMC3* | 0 | -0.6 | -1.8 |
| *ITGB8* | 0 | -0.7 | -1.9 |
| *SHC2* | 0 | -0.7 | -2.0 |
| *ITGA11* | 0 | -0.8 | -2.0 |
| *CD36* | 0 | -0.8 | -2.0 |
| *IQGAP2* | 0 | -0.2 | -2.5 |
| *MMP11* | 0 | -0.5 | -2.8 |
| *ITGA8* | 0 | -1.3 | -2.9 |
| *KDR* | 0 | -1.3 | -3.5 |

